## Supplementary figures for "Neuroprotective role of Hippo signaling by microtubule stability control in *C. elegans*"

Lee_Supplemental_Fig 1

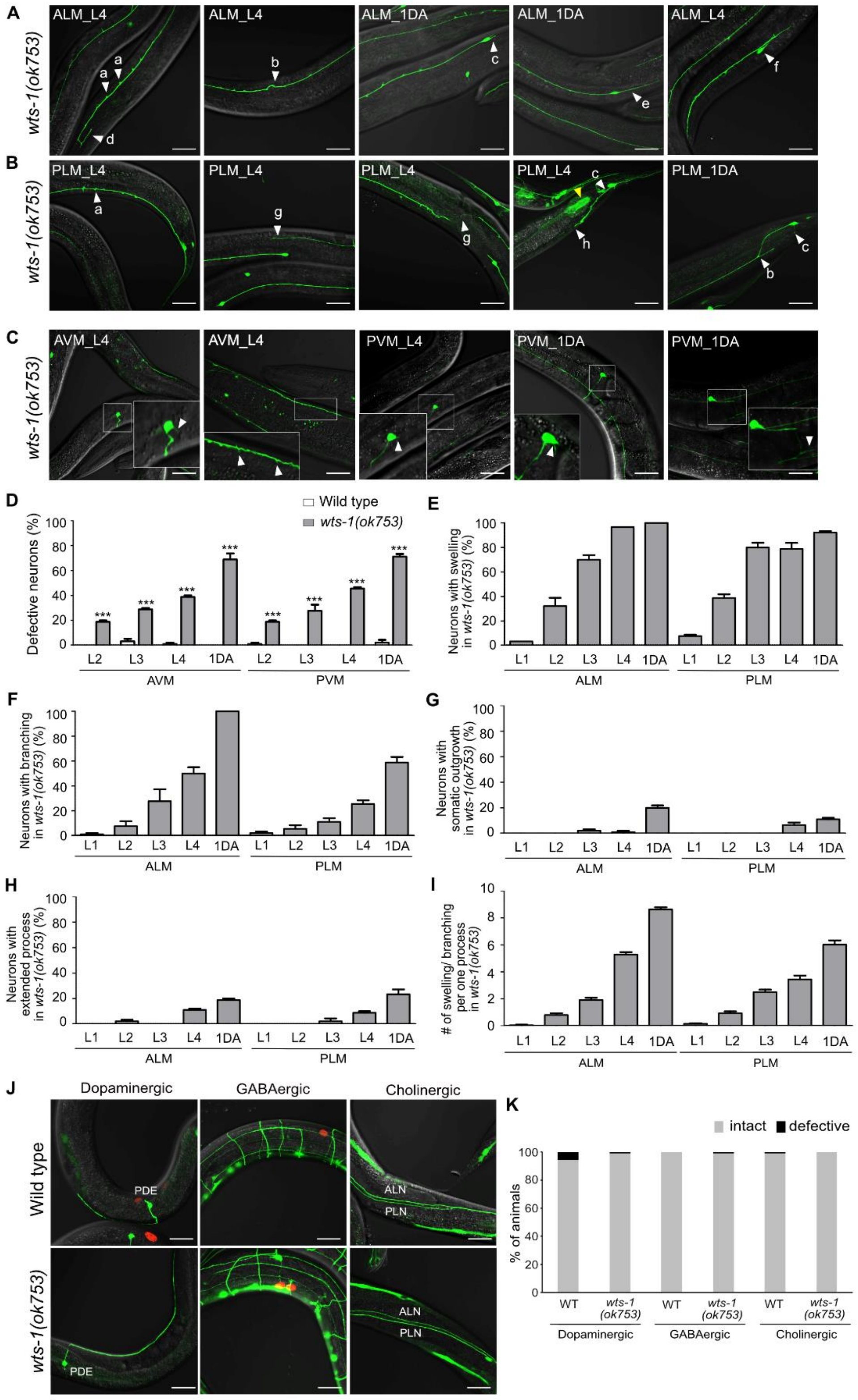

Lee_Supplemental_Fig 2

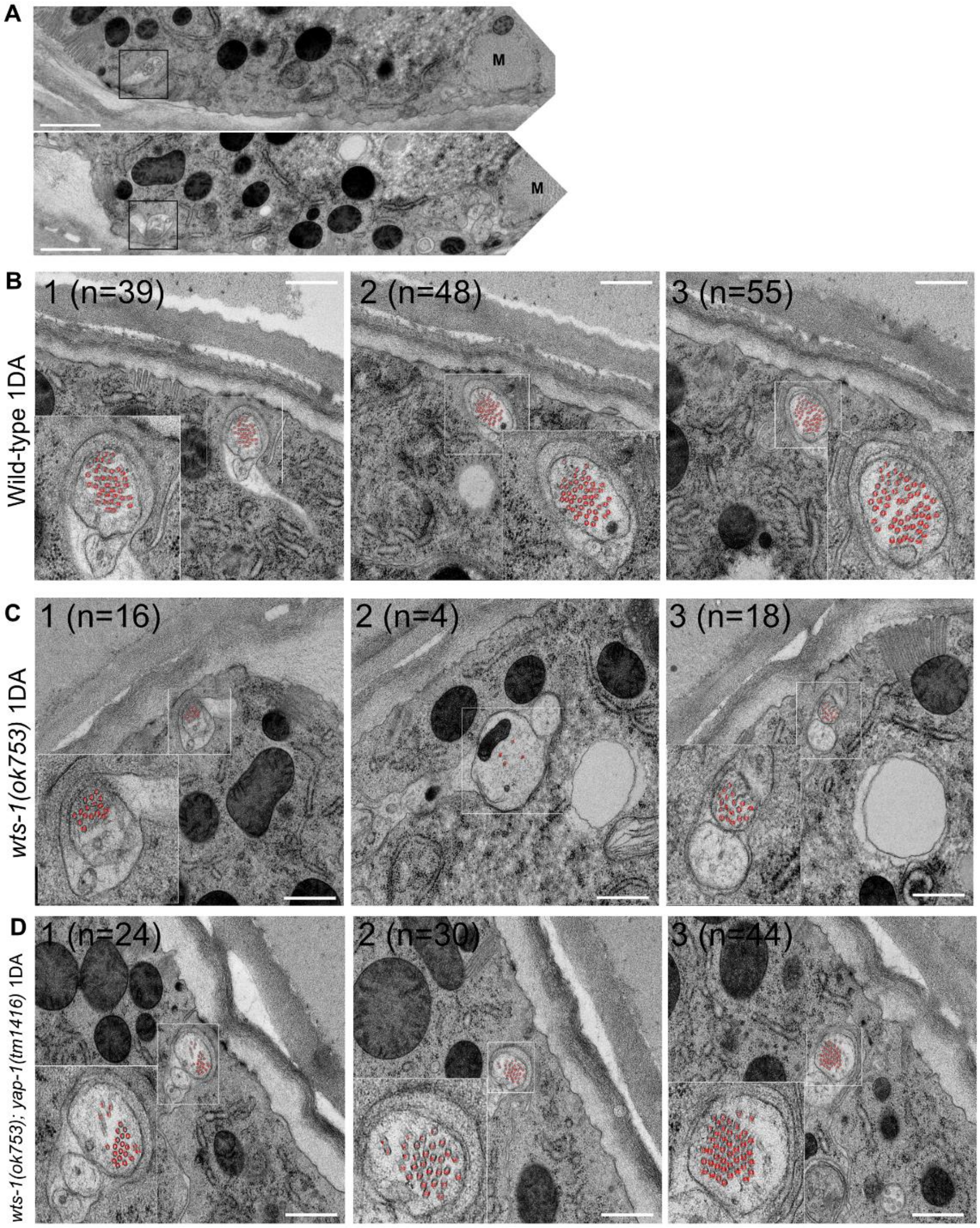

**Lee_Supplemental_Fig 3**

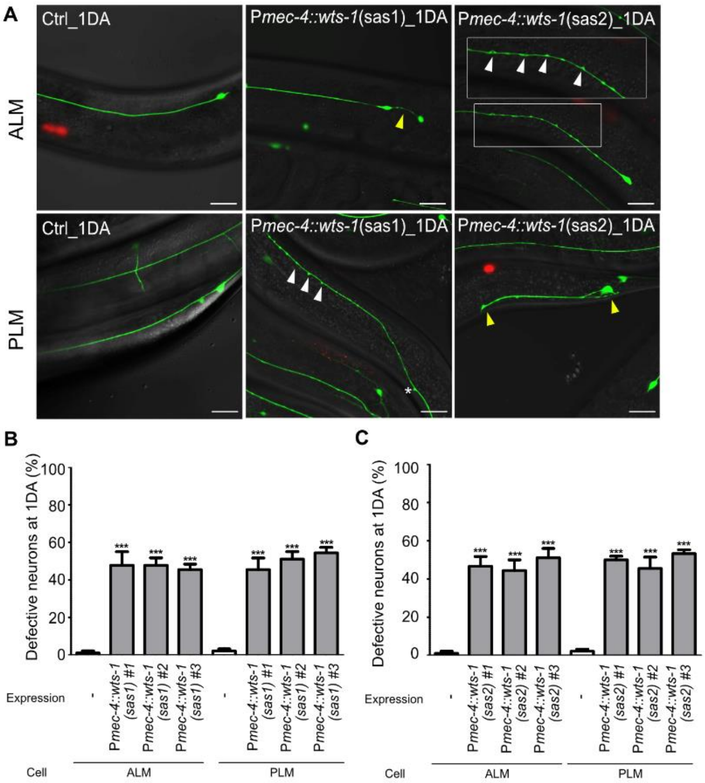

**Lee_Supplemental_Fig 4**

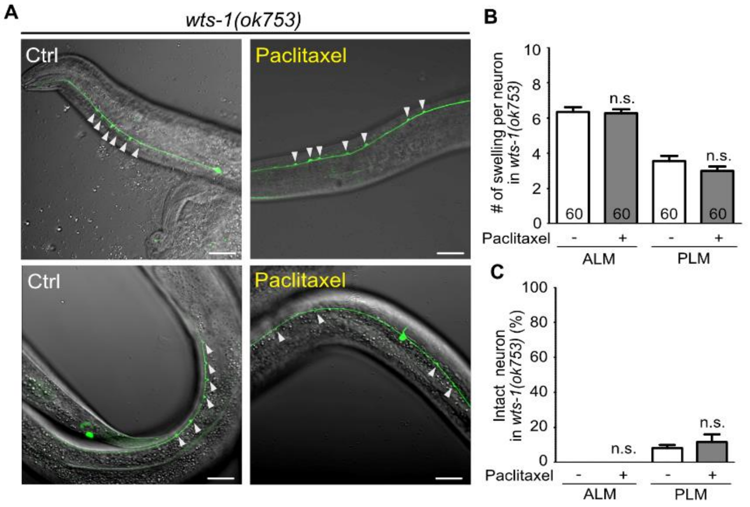

Lee_Supplemental_Fig 5

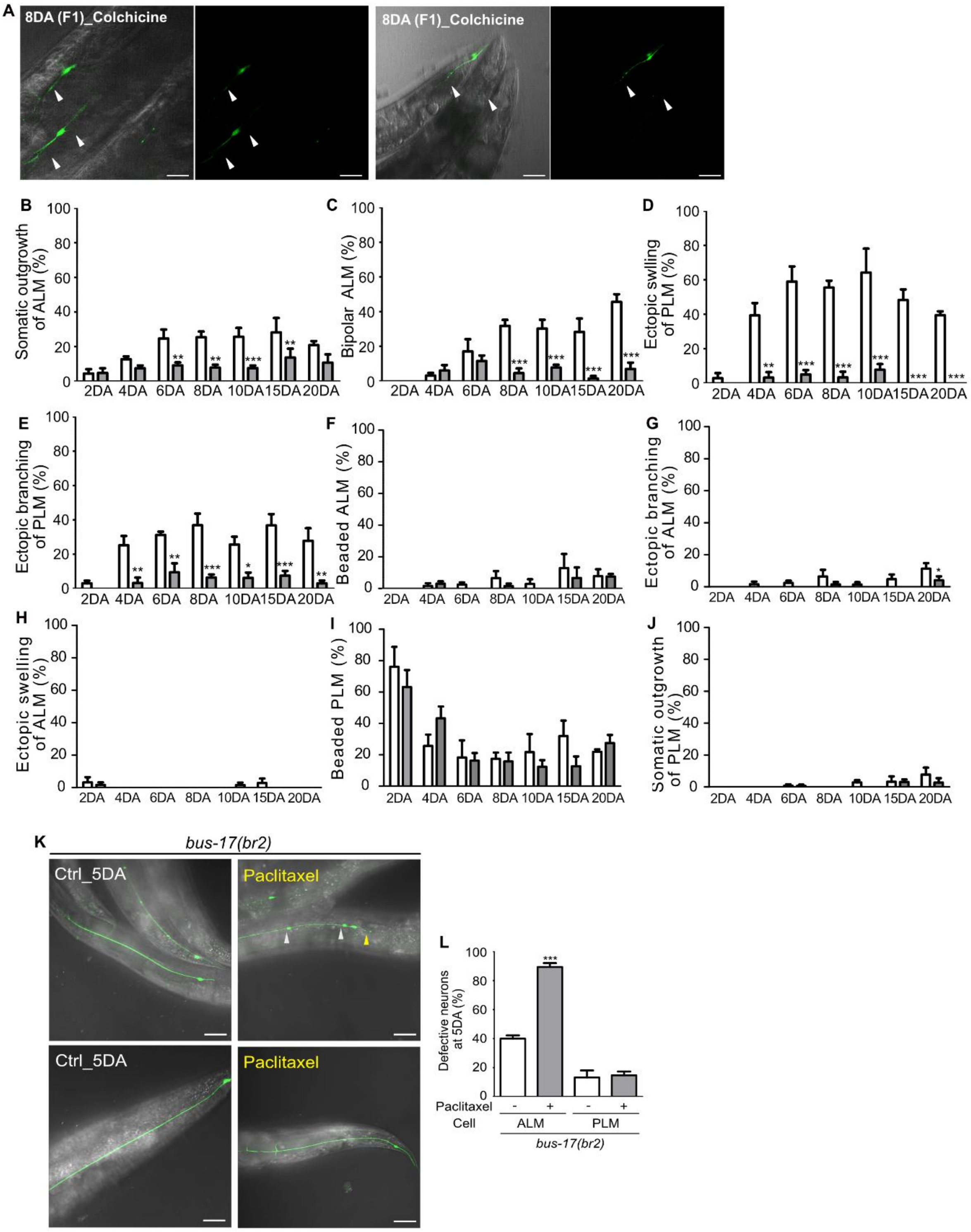

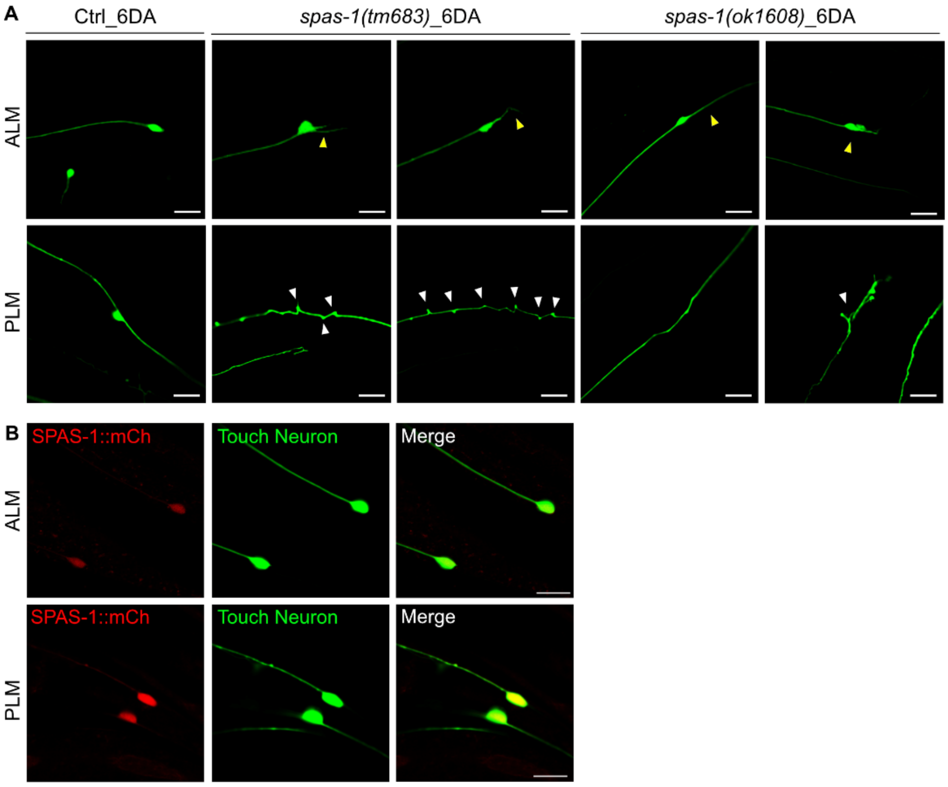
A
Ctrl_6DA
spas-1(tm683)_6DA
spas-1(ok1608)_6DA
ALM
PLM
B
SPAS-1::mCh
Touch Neuron
Merge
ALM
SPAS-1::mCh
Touch Neuron
Merge
PLM

Lee_Supplemental_Fig 7

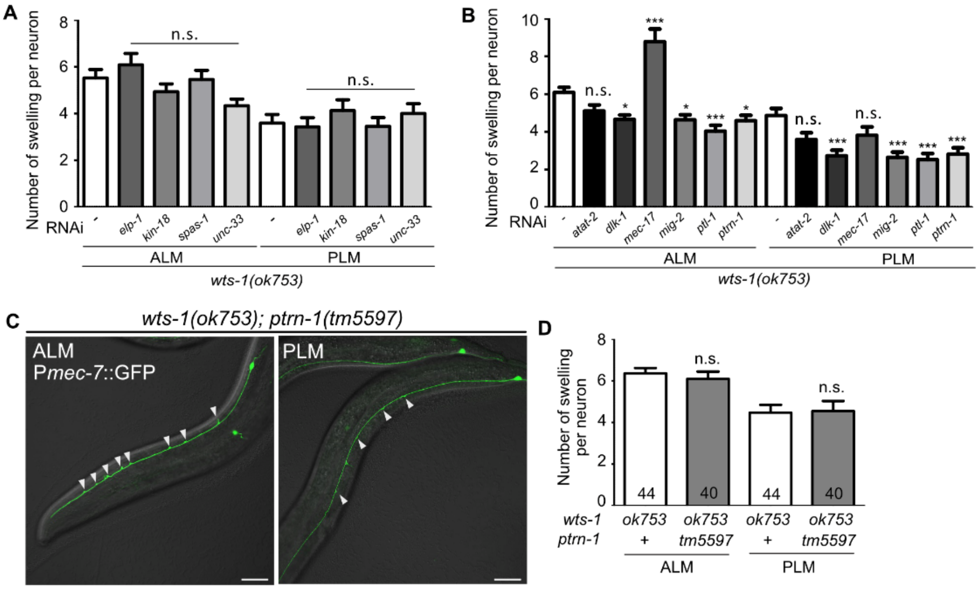
